# Never judge a bird by its feathers: genetics unveils the true target of trafficking in morphologically similar *Copsychus* species

**DOI:** 10.1101/2023.04.14.536889

**Authors:** Emily L. Cavill, Germán Hernández-Alonso, Rob Ogden, M. Thomas. P. Gilbert

## Abstract

Trends in wildlife crime continue to rise and contribute to the ongoing decline of our planet’s biodiversity. We applied a genetic approach to ascertain the geographic origin of a suspected Seychelles Magpie-robin confiscated in Singapore during international transit, and to confirm illegal trade of this species. However, mitochondrial analyses revealed this individual to be a subspecies of Oriental Magpie-robin, endemic to the island of Borneo, and bearing similar morphology to the Seychelles species. We thus consider the implications of correct species identification on captive care and repatriation in cases of wildlife confiscation, and emphasise the value of using genetics in wildlife crime investigation.

## Introduction

The World Wildlife Fund recognises wildlife crime, such as the illegal poaching or trafficking of flora and fauna, as the second biggest threat to threatened species and their ecosystems (WWF, 2020), and several recent studies (Morton *et al*., 2021; Scheffers *et al*., 2019) have documented that wildlife trade is accelerating the rate of species extinction. While there are wider implications of both the *legal* wildlife trade (such as poor regulation, animal welfare issues, and forgeries as discussed in Wyatt *et al*., 2022) and *illegal* wildlife trade, given that the latter is totally unregulated, its consequences are the most severe. For example, while it has been estimated that the illegal wildlife trade is a multi-billion-dollar industry (Andersson *et al*., 2021), the local economies in which it occurs suffer due to declines in wildlife tourism, loss of work, food, resources, and taxes (Cardoso *et al*., 2021). A further negative consequence is that poaching directly reduces biodiversity while also disturbing the delicate balance of ecosystems through the removal of native species and the introduction of alien species. And lastly, international wildlife trade can remove geographic barriers between species which when broken through anthropogenic movement can facilitate introgression or hybridisation events (reviewed in Adavoudi & Pilot, 2022), may introduce novel diseases to which native species have no resistance, and can lead to increases in outbreaks of zoonotic diseases in the human population (Jones *et al*., 2008; Rush *et al*., 2021).

An assessment of data including records, databases, and reports compiled through the IUCN (International Union for Conservation of Nature) and CITES (Convention on International Trade in Endangered Species of Wild Fauna and Flora) has identified birds as the most commonly traded of the terrestrial vertebrates that are listed in the Red List ‘threatened’ (Critically Endangered, Endangered, Vulnerable, Data Deficient) categories (Scheffers *et al*., 2019). In general, trafficking of wild animals is driven either by a species’ popularity (where more exotic and rare species are sought after), or because customers are simply willing to pay a premium for wild-caught (as opposed to cage-bred) birds (Ribeiro *et al*., 2019; Chng *et al*., 2021).

The research presented here arose when a suspected Seychelles Magpie-robin, *C. sechellarum* (SMR), was confiscated live in transit to Singapore in 2019, and subsequently cared for by Singapore’s bird park, Jurong. The Magpie-robin and Shama (MRS) taxonomic/species complex is often referred to under the genus *Copsychus*. While Lim *et al*. (2010) proposed three genera be named within the complex (*Copsychus, Kittacincla, Trichixos*), Voelker *et al*. (2014) later proposed a fourth (*Saxicoloides*). These differing names add to the ongoing taxonomic confusion concerning this group, especially as they are not universally adopted in the discussion of this complex. While we recognise these synonymous names, for the purpose of this paper we solely use the genus name *Copsychus* when referring to the multiple species within this group. The *Copsychus* genus is widespread across South/Southeast Asia, and species are also found in two countries of the Western Indian Ocean, Madagascar and Seychelles. While there is little concrete evidence of trade in the African species, the Asian Magpie-robins and Shamas are highly targeted and heavily traded locally and internationally (Burivalova *et al*., 2017; Chng *et al*., 2018; Leupen *et al*., 2018). Due to this, many species and subspecies within the complex are suffering drastic and, in some cases, irreparable population declines. The Seychelles Magpie-robin is an IUCN red-listed ‘Endangered’ species, endemic to Seychelles, and holds high social importance. There are fewer than 500 individuals existing in Seychelles, and the species, as with most birds in Seychelles, is protected under the national legislation of the Wild Animals and Birds Protection Act (WABPA, Seychelles). Through recent online monitoring, the Seychelles Magpie-robin has potentially been identified as a subject of the illegal pet trade; bred in cages in Java, Indonesia, and sold under the trade name ‘Kacer Wulung’ (S. Bruslund, *pers. comm*.). However, confirming these are the Seychelles species requires more on-the-ground investigation. Thus, the prospect of obtaining genetic material of a Seychelles Magpie-robin being trafficked internationally offered hope that we may be able to provide incontrovertible evidence of this illegal activity. Our main aim was to find the local source of the poaching through comparison to our existing, extensive genomic SMR reference dataset, to improve protective measures in place and offer justification for international protection, i.e. applying for a CITES appendix.

The use of genetic approaches as a tool for species identification is increasingly applied in the forensic investigation of wildlife crime and is particularly relevant for species where identification based on morphology allows room for ambiguity. Three groups within the *Copsychus* genus are characterised by the melanistic black-bellied *and* black-tailed phenotype as observed in the Singapore’s bird park Magpie-robin (herein referred to as JMR) being investigated in this study: both sexes of the Seychelles Magpie-robin (*Copsychus sechellarum*, Figure 1A) endemic to Seychelles; the males of a subspecies of Oriental Magpie-robin (*Copsychus saularis adamsi*, Figure 1B) largely restricted to Sabah, Borneo; and the males of the black-bellied Madagascar Magpie-robin (the nominate race of *Copsychus albospecularis*, Figure 1C) endemic to Madagascar. As the JMR is morphologically akin to these groups, genetic assessment was imperative to accurately assign species to this individual to be able to further explore the implications of the trafficking event. We re-sequenced the whole genome of the JMR and extracted the mitochondrial genome for phylogenetic analyses to test our belief that this individual was an SMR.

**Figure 1.**
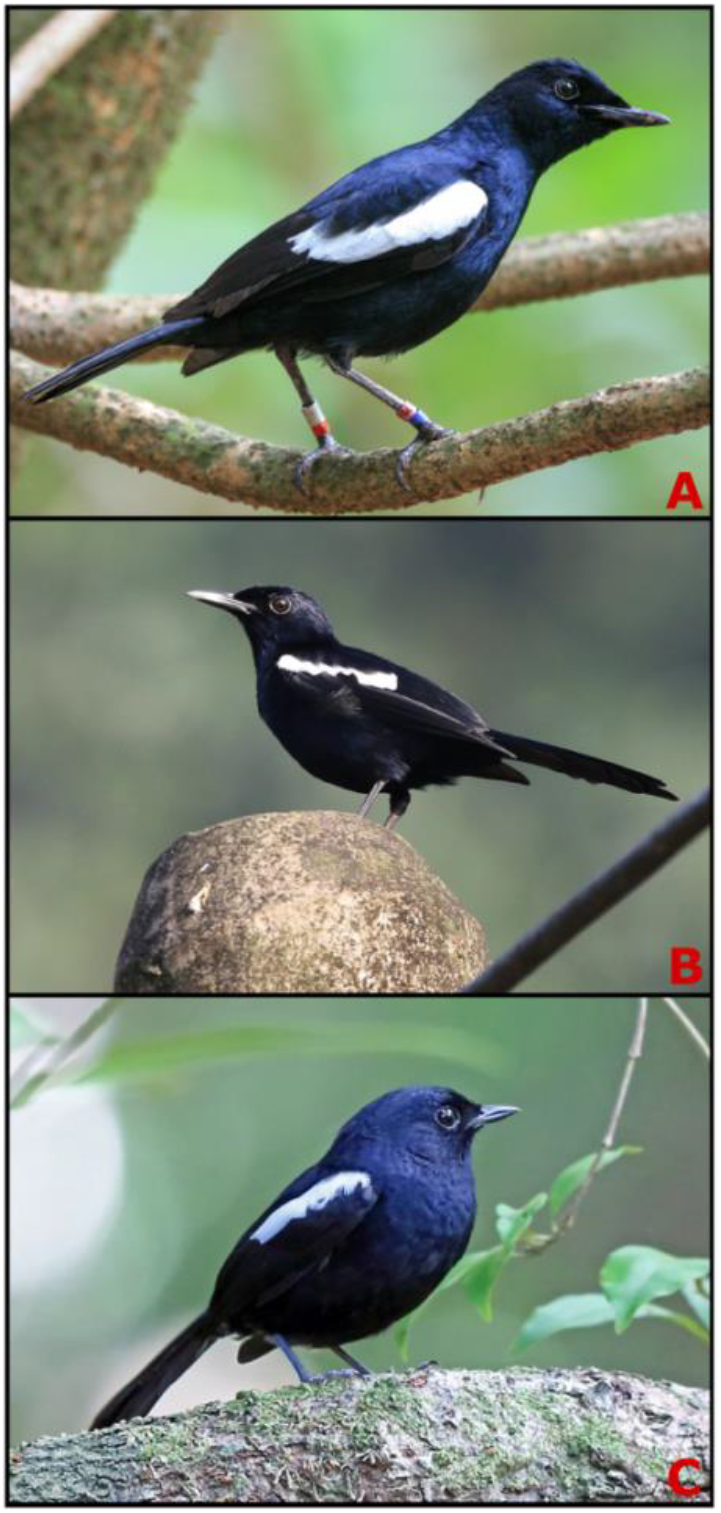
Photographs demonstrating the morphological similarities between the three totally melanistic Copsychus species. A) Photograph of Seychelles Magpie-robin taken by Oscar Campbell. B) Photograph of Oriental Magpie-robin adamsi taken by Richard Greenhalgh. C) Photograph of Madagascar Magpie-robin taken by Nigel Voaden. All photographs provided by the Cornell Lab of Ornithology under a non-commercial Media Licence Agreement.

## Methods & materials

### Sample collection and lab processing

A total volume of 40μL (10% of body weight) of blood was drawn from the jugular vein of the confiscated JMR residing at Singapore’s bird park. The sample was collected by a resident vet and stored in 1.5mL of lysis buffer. The blood sample was exported from Singapore under permit number NP 1.35.1.1.3/070722 to the Globe Institute, University of Copenhagen, Denmark, for research under permission from the Ministeriet for Fødevarer, Landbrug og Fiskeri (J.nr. 2022-12-7186-02006).

DNA was extracted following the DNeasy Blood & Tissue protocol (Qiagen), and subsequently fragmented to ∼350 bp (Covaris LE220-plus, 25 μL input DNA with settings 110s, peak power 200, duty % factor 25, cycles per burst 50, average power 50). A BGISeq compatible shotgun sequencing library was built on the fragmented DNA following Kapp *et al*. (2022), as modified for the BGIseq platform following (Van Grouw *et al*., in review). The library was amplified through a single 50 μL reaction using 5 μL library template, 1x AmpliTaq Gold buffer, 2.5 mM MgCl2, 0.2 mM dNTP, 0.2 μM of both forward and reverse primer, and 0.1 U/μL AmpliTaq Gold polymerase, (with thermal cycling conditions of 95°C/12 m, followed by 12 cycles of 95°C for 20 s, 60°C 30 s, 72°C 40 s, followed by 72°C for 5 m), and the amplified library was purified with a Hi-Prep PCR clean-up system (MagBio). Sequencing was performed on the BGI DNBseq4000 platform using 150PE reads by BGI’s commercial service.

### Taxonomic assignment

Given the morphological challenges surrounding species identification, outlined above, a phylogenetic approach was used to infer the genetic affinity of the JMR sample to in-situ SMR and other species of the genus.

We generated mitochondrial genome bam files using PALEOMIX v1.3.6 BAM pipeline (Schubert *et al*., 2014). First, adapters were trimmed using AdapterRemoval v.2.3.3 (Schubert *et al*., 2016). Trimmed reads for the JMR were mapped to reference mitochondrial genomes for SMR (accession number MN356447) and OMR (accession number KU058637) with BWA v0.7.1 mem algorithm (Li & Durbin, 2009) using default parameters. Picard MarkDuplicates (Broad Institute, 2019) was used to identify and remove PCR duplicates. Additionally, genomic sequences of five SMR, one per in-situ island (retrievable through Bioproject number PRJNA722144) were included in analyses to expand the SMR comparative dataset and potentially identify the origin island of the sample, if it was an SMR. These data were mapped to the SMR reference mitogenome under the same conditions described above. Additionally, publically available DNA sequences for the entire NADH dehydrogenase subunit 2 (ND2) gene for 74 *Copsychus* samples representing 20 taxa were downloaded from NCBI. A comprehensive list of accession numbers for all data used in this study are detailed in Table 1. The consensus command in samtools v1.16 (Danecek *et al*., 2021) was used to convert the bam files generated for the JMR and SMR to consensus fasta sequences for the mitochondrial genome, not allowing insertions or deletions in the final sequence. The mitogenome fasta sequence for *Cercotrichas coryphaeus* (Karoo Scrub-Robin (KSR), accession number MN356422) was downloaded to use as an outgroup for all analyses. We then extracted the entire nucleotide sequence for the ND2 mitochondrial gene from the whole mitogenome fasta files (JMR, SMR, OMR, and KSR) using the getfasta command in bedtools v2.9 (Quinlan & Hall, 2010), supplying a coordinate file in bed format.

**Table 1:**
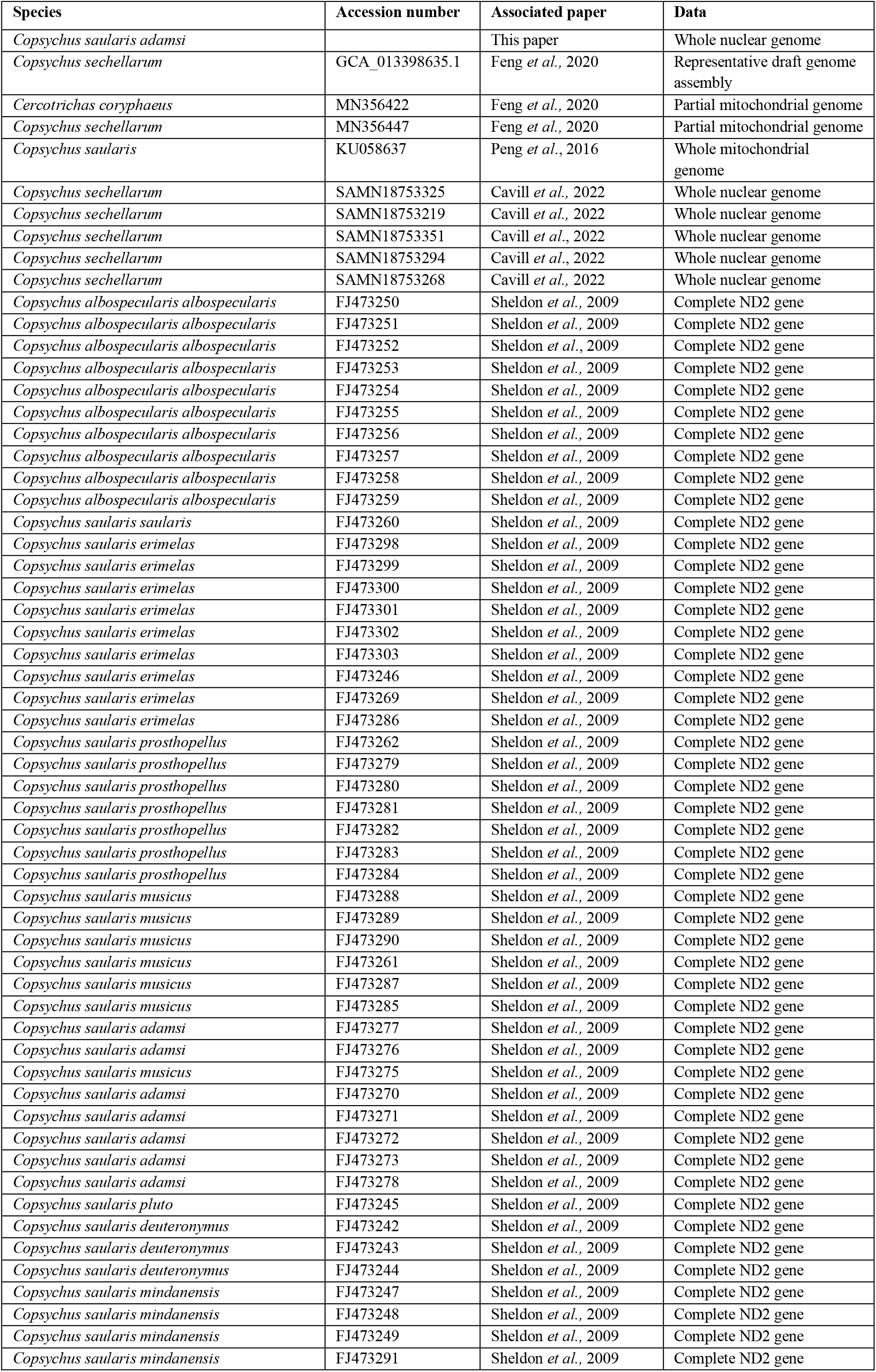
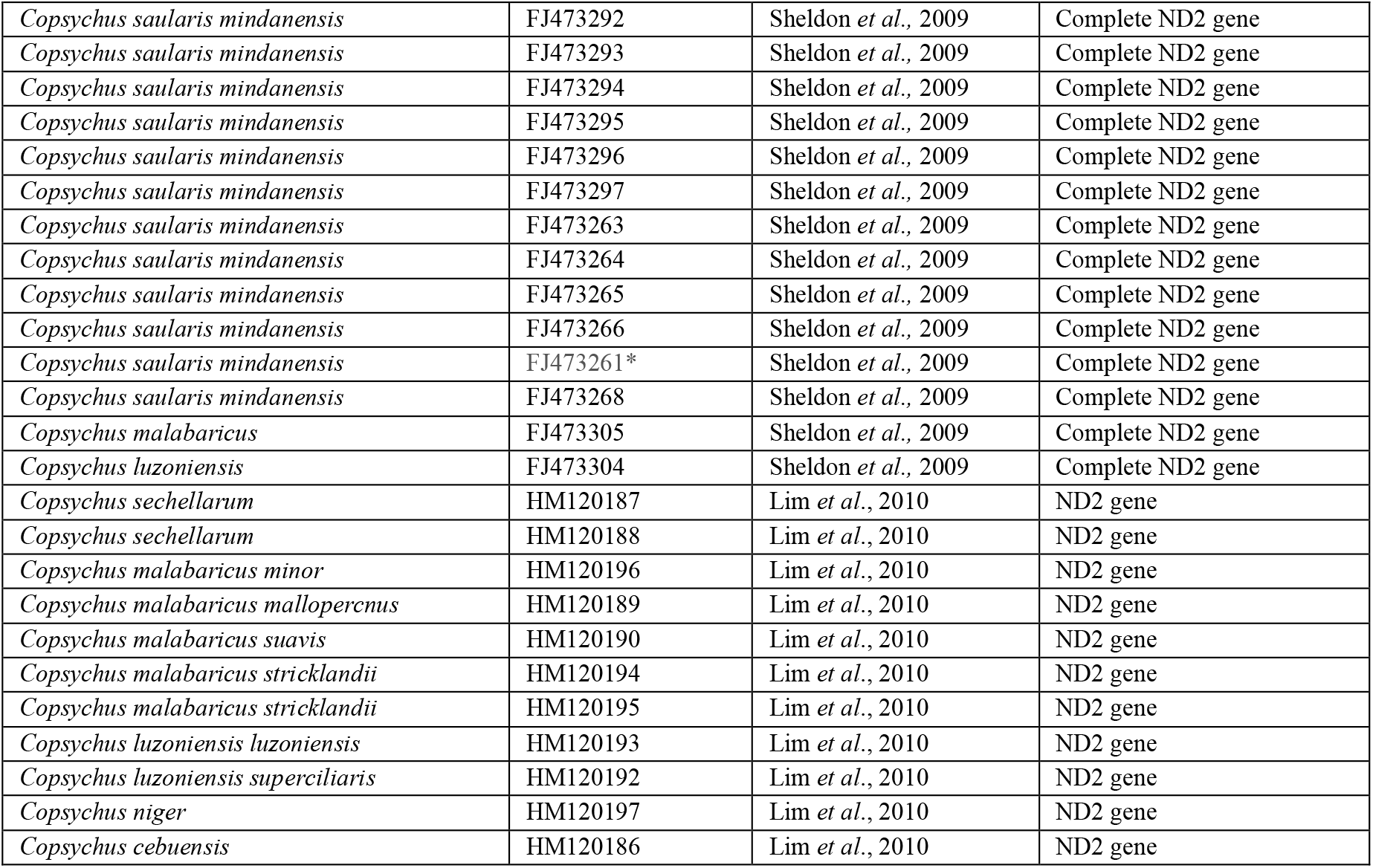
Table containing details of all samples used in this study. Species name, accession number, associated paper and data type used for each sample included in the analyses in this research. ND2 = NADH dehydrogenase subunit 2 (ND2) gene. FJ473261* listed in its original paper as FJ473260 but it was a duplicate no. and wrong species, we searched for the sample details in the database and retrieved the included number.

We generated two trees: the first, a whole mitochondrial genome phylogeny for the JMR, and the high-quality draft reference assembly mitochondrial sequences of the SMR and OMR, and KSR, using the KSR as an outgroup; and the second using only the ND2 genes for all 83 samples included in this study. First, we created a multiple sequence alignment using mafft v7.505 (Katoh & Standley, 2013). We then used modeltest-ng v0.1.7 (Darriba *et al*., 2019) to select the best substitution model. Subsequently, 20 repetitions of independent maximum likelihood trees were created with raxml-ng v1.1.0 (Stamatakis, 2014) using substitution model GTR+I+G4 and 1000 bootstrapped estimates were generated. The estimated phylogenetic trees were visualised using the Interactive Tree of Life (iTOL) v5 (Letunic & Bork 2019).

## Results

From the whole genome resequencing data of the JMR, 6850 reads mapped to the SMR and 9355 reads mapped to the OMR reference genomes, yielding a mitochondrial genome coverage of 75x and 105x respectively. The whole mitochondrial genome analysis revealed the JMR to be most closely related to the OMR (Figure 2). In this phylogeny, the SMR presents as a sister group to the OMR, consistent with findings in Lim *et al*. (2010). Subsequently, we only present the results generated using the OMR-aligned mitochondrial genome. Further analysis limited to the NADH dehydrogenase subunit 2 (ND2) gene, to allow the inclusion of a large number of reference samples, indicated that the JMR is the *adamsi* subspecies of the OMR (Figure 3). Taking into consideration the melanistic morphology of the JMR, the sex of the bird, the divergence estimates of these three melanistic species (Lim *et al*., 2010), and the high branch support value obtained for the whole mitogenome phylogeny (0.94) and ND2 phylogeny (0.96-1.0) it is extremely likely that the subspecies of this individual has been correctly assigned.

**Figure 2.**
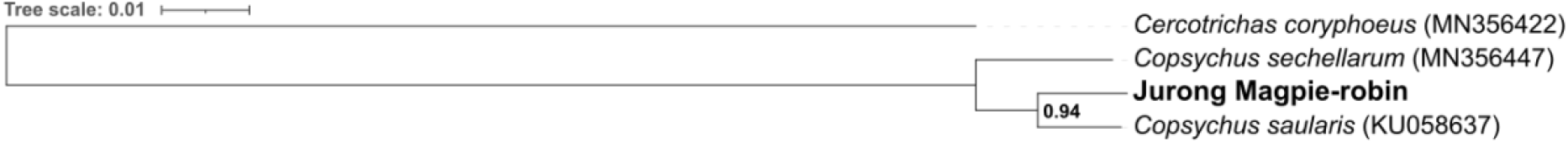
Mitochondrial genome maximum likelihood phylogenetic tree identifying the species of the JMR. Phylogeny created using whole mitochondrial genomes of the Seychelles Magpie-robin (Copsychus Sechellarum), the Oriental Magpie-robin (Copsychus saularis), and the Singapore bird park (Jurong) Magpie-robin (highlighted in bold). The Karoo Scrub-robin (Cercotrichas coryphoeus) was used as the outgroup. JMR clearly groups with the Oriental Magpie-robin, supported with a high bootstrap value, while the Seychelles Magpie-robin forms a sister group.

**Figure 3.**
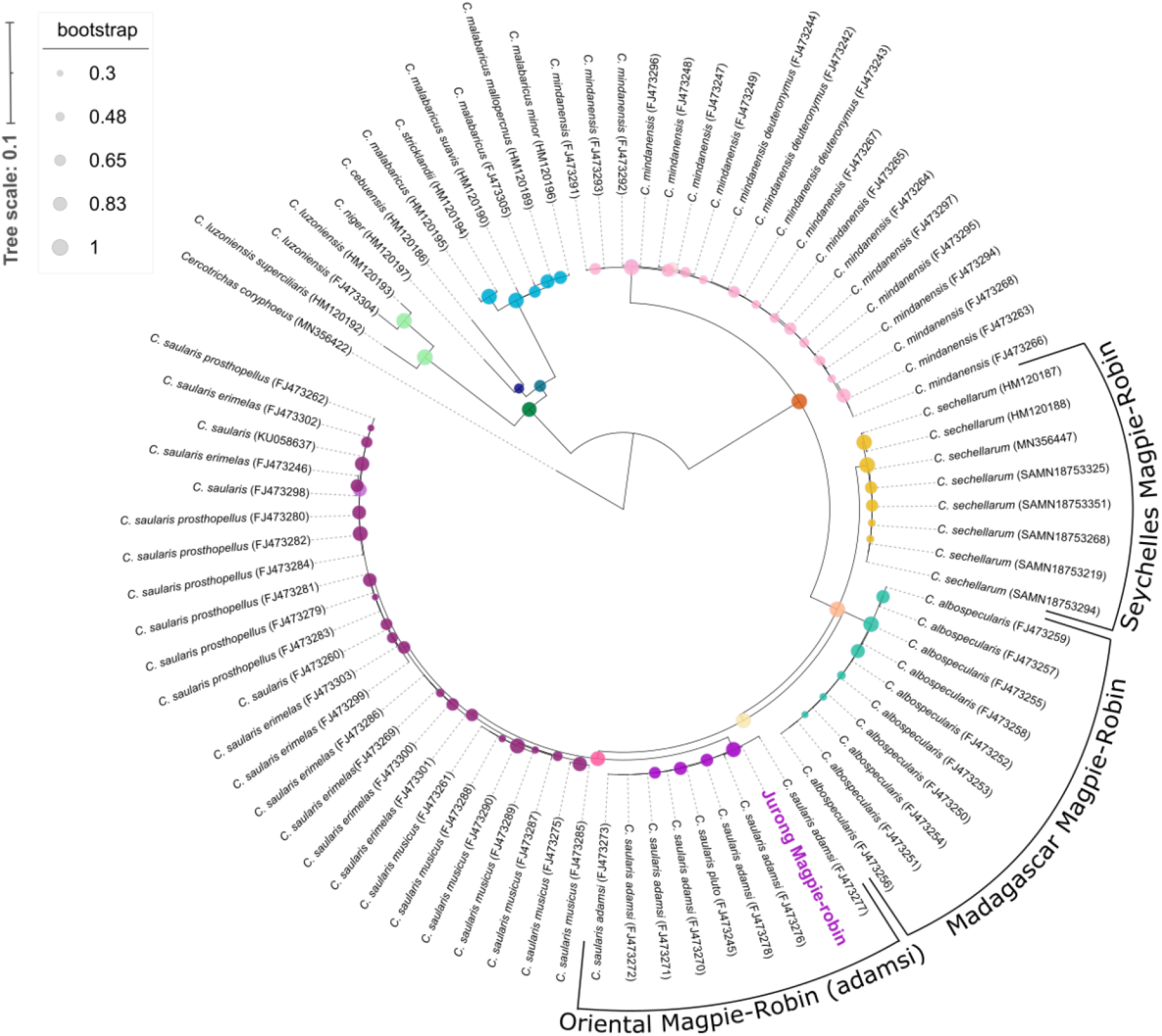
Maximum likelihood phylogenetic tree for the ND2 gene of Copsychus species. Bootstrap support defined in the legend, reported from 0.3-1.0. Groups with melanistic phenotypes are labelled with common names. Current nomenclature used in individual identifying labels (i.e., C. mindanensis and C. stricklandii which are now recognised as full species). Karoo Scrub-robin (Cercotrichas coryphaeus) presented with full latinised name, all ‘C.’ abbreviations are Copsychus species. The position of the Singapore bird park (Jurong) Magpie-robin, highlighted in purple, sits within the adamsi subspecies of Oriental Magpie-robin with high bootstrap support.

## Discussion

This research originated in light of the trafficking of a suspected Seychelles Magpie-robin into Singapore. As previously stated, the males of the *adamsi* subspecies of the Oriental Magpie-robin share the melanistic phenotype with the males of the nominate race of the Madagascar Magpie-robin, and both sexes of the Seychelles Magpie-robin. These shared morphological characteristics evidently led to the misidentification of this individual at the point of confiscation. Correct identification of a species can determine the severity of the crime committed based on legislation surrounding poaching or trafficking of a species, for example if the species is CITES listed, and thus inform the prosecution. Morphology is often the first step of species identification in cases of wildlife crime, and further genetic testing may not be necessary in very obvious cases (e.g. tiger skin, elephant tusks, pangolin scales). However, genetic testing can also be implemented in these cases to determine the geographical origins of animals or their parts to identify poaching hotspots (e.g. Wasser *et al*., 2008). Additionally, in cases where live animals are confiscated and cared for in an ex-situ environment (following one of the main CITES guidelines for live animal seizures, IUCN 2019), incorrect species identification has ramifications for care in captivity and inclusion in breeding and reintroduction programs. Fortunately, we have now been able to remedy this in the case of the JMR. Further complications of misidentification occur if animals are returned to their assumed originating country (additional CITES guideline) as releasing animals into non-native habitats, or without understanding of genetic population structure can be detrimental to in-situ populations. Given that the SMR are burdened with limited genetic diversity (Cavill *et al*., 2022), repatriation of trafficked birds could be vital for the species survival. If the JMR had been ‘returned’ to Seychelles, it would have exposed the Seychelles species to novel pathogens and potentially resulted in genetic introgression. Hence, the case of the JMR illustrates the value of genetics in the investigation and subsequent management of illegal trade in live animals, especially the management of confiscated livestock.

There is long-standing taxonomical ambiguity within the *Copsychus* genus due to its paraphyletic nature and ongoing resolution of lineages. Within the MRS complex, two large groups have proved the most difficult to disentangle: *malabaricus* and *saularis*. We have already witnessed how the progression of genetics has been used to identify full species that were previously conspecific of *malabaricus* or *saularis* (namely *C. mindanensis*, Sheldon *et al*., 2009, and *C. superciliaris*, Lim *et al* 2010). More recently, through the generation of whole-genome re-sequencing data, Wu *et al*. (2022) identified four full species within the existing *C. malabaricus* group. This has also created an extensive genomic database against which to test future cases of illegal poaching/trafficking of this recently CITES II listed species (CITES, 2023). However, there is still much uncertainty within *C. saularis*, and Lee *et al*. (2016) cautions there may be full species within this group. Taxonomic complexities can be understood and ultimately resolved with the accruement of knowledge, increased availability of genetic resources, and advancements in technology. Thus, the *saularis* species will undoubtedly benefit from the implementation of contemporary genomic methods to identify any distinct lineages. This will not only enable assessment for appropriate conservation units but will facilitate species assignment in future trafficking incidents.

The OMR is a species under which 12 subspecies have been recognised (Gill & Donsker, 2014). The latest IUCN assessment (2020) considers the OMR of ‘Least Concern’ as it does not meet the ‘vulnerable to threat of extinction’ threshold. While it is true that the OMR as one species complex has a large geographical range across Asia, spanning from China to Indonesia to Pakistan, with a notably large and ‘stable’ population, this encompassing conservation status classification neglects the ‘pockets’ of smaller, geographically, and potentially genetically isolated groups within this species that may be evolutionarily distinct and warrant different conservation status and different conservation strategies. Analyses of the market trade (Harris, 2015) and other assessments (Lee *et al*., 2016; Chng *et al*., 2021) indicate many groups within this species complex are in decline, with some possibly close to extinction in the wild. The *adamsi* subspecies of OMR, to which we have established the JMR belongs, is now regarded as rare in the wild (Lim *et al*., 2017) and may soon be lost without conservation efforts. There is indication of genetic swamping of the black-bellied *adamsi* and *pluto* on Borneo by the white-bellied *musicus* (Sheldon, 2009). However, as *adamsi* have a restricted endemic range in Northeast Borneo, but have been identified being sold in pet shops in Singapore (Eaton *et al*., 2017), and the JMR was confiscated in transit from Malaysia to Singapore (NParks, *pers. comm*.), there is also compelling evidence of international trade of this group. *Copsychus stricklandii, malabaricus*, and *saularis* are all legally protected in Sabah through the Wildlife Conservation Enactment (as discussed in Chng *et al*., 2021), and thus it is illegal to catch and remove *C. s. adamsi* from the wild. However, a lack of compliance and effective law enforcement suggests that local protection alone is insufficient (Chng & Eaton, 2016), and this is further demonstrated by the case of the JMR. To give some insight into the extent of trafficking of the OMR, over 17,000 OMR were seized being smuggled out of Malaysia in 2020 alone (Lee *et al*., 2016). Chng *et al*. (2021) have proposed that the OMR be added to additional existing national wildlife protection laws, and ultimately the species should be granted a CITES listing for international protection.

Although physical bird markets are still rife throughout Asia, many traders now take advantage of access to poorly regulated online marketplaces (Siriwat & Nijman, 2020; Okarda *et al*., 2022). Investigations which monitored online wildlife trade in Malaysia found the OMR were the 9th most traded animals on Facebook, a prominent online market for wildlife trade (Chng *et al*., 2021). The SMR is another species which can be found, throughout the Facebook online marketplace, advertised for sale in South East Asia (Figure 4). While the logistics involved in international smuggling of the SMR seem complicated and perhaps unrealistic, and considering that other, geographically closer, melanistic Magpie-robins can be easily mistaken for the SMR, it may transpire that the morphologically similar *adamsi* is deceptively traded under the highly endangered, incredibly rare and exclusive SMR species to drastically increase the price to collectors, although *adamsi* is known to be traded under a different name. This isolated case of misidentification of the JMR should not be used to refute findings, made through online monitoring, that the SMR are subjected to international trafficking. Given that the SMR is endemic to the Seychelles, if birds of this species are being sold in Indonesia there has unquestionably been illegal poaching from the wild. We therefore advocate for further investigation into the SMR trade in Indonesia, through wildlife forensic research, to ascertain if this species is a target of illegal wildlife trade.

**Figure 4.**
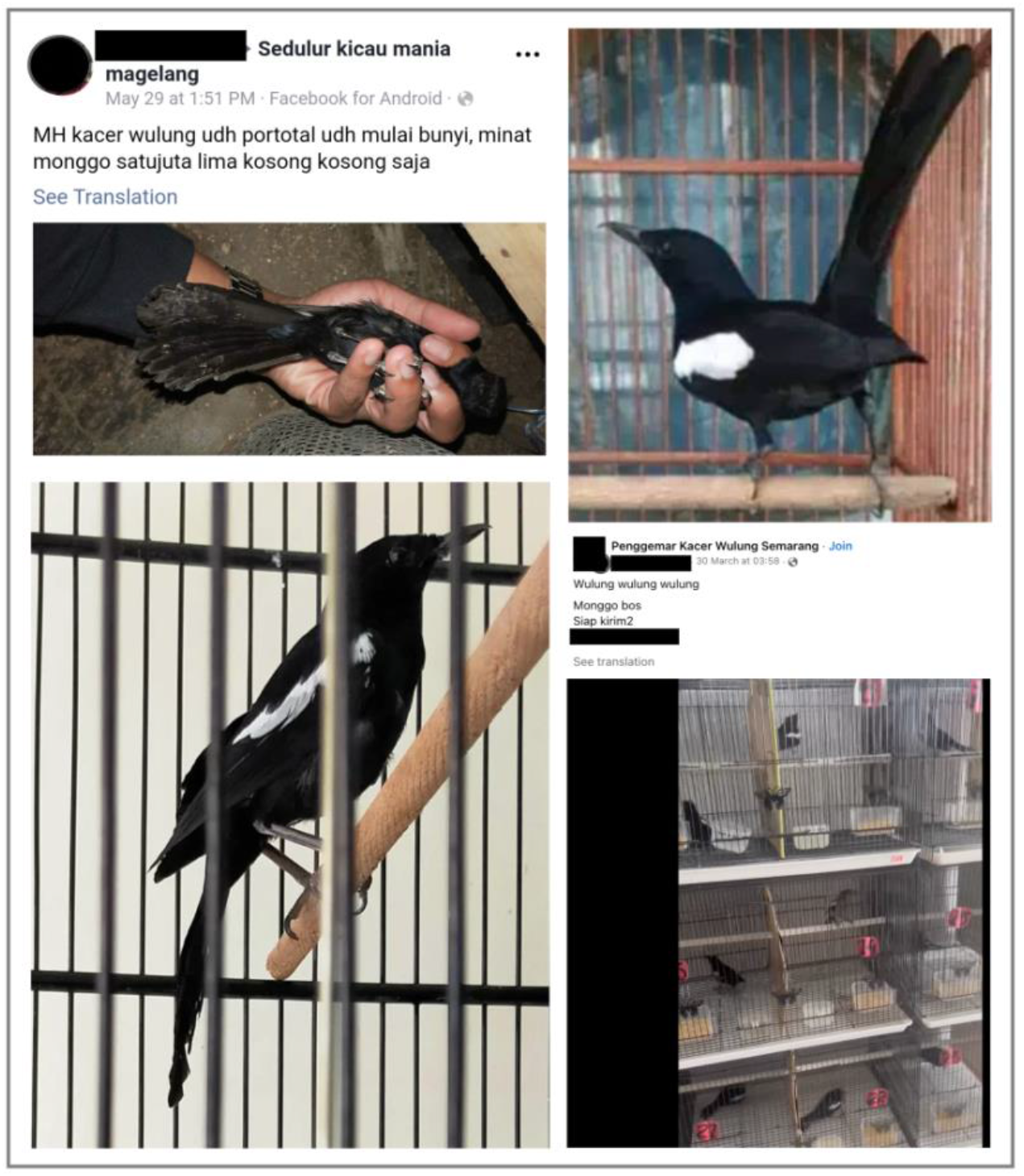
A panel of images of SMR advertised for sale under the name ‘Kacer Wulung’, retrieved through Facebook. Images of claimed-to-be Seychelles Magpie-robins collected from Facebook in 2019 (left) and 2023 (right), the birds in which clearly possess the black-bellied and black-tailed melanistic phenotype associated with this species. These images are freely viewable through a search on Facebook without requiring membership to any groups, however any identifying details of the original poster (photos, names, and phone numbers) have been covered.

## Conclusions

Although we are already aware of the high trafficking rates of the OMR in Asia, through genetics we have now been able to identify a case of trafficking of a rare-in-the-wild and legally protected OMR subspecies and communicate this finding to the relevant authorities. We strongly reiterate the importance of species identification through genetic analysis in cases of wildlife crime, particularly pertaining to the subsequent management of confiscated live animals. Finally, there is a high-quality genomic database available both for the entire range of in-situ SMR and the OMR(*adamsi*), and we recommend the investigation of the detected illegal SMR trade in Indonesia be extended with the implementation of genetic tools in the species assignment of these caged birds.

## List of abbreviations

CITES: Convention on International Trade in Endangered Species of Wild Fauna and
Flora IUCN: International Union for Conservation of Nature
JMR: Singapore’s bird park (Jurong)
Magpie-robin KSR: Karoo Scrub-robin
MRS: Magpie-robin and Shama complex
NParks: National Parks Board
OMR: Oriental Magpie-robin
SMR: Seychelles Magpie Robin
WWF: World Wildlife Fund

## Acknowledgements

We are most thankful to the veterinary staff at Singapore’s Bird Park, Jurong, for their cooperation and expertise in sampling the Magpie-robin. We extend these thanks to Jessica Lee of Mandai Nature for facilitating sample collection and for her helpful feedback on the final draft of this manuscript. We are grateful to Singapore National Parks Board (NParks) for their cooperation, especially Gerald Neo for organising the export permit. We also thank Mandy Bolt Botnen and Daniel Bilyeli Øksnebjerg at the Globe Institute, University of Copenhagen, Denmark, for facilitating permits for the import of avian samples into Denmark. We thank Simon Bruslund of Copenhagen Zoo for his valuable knowledge of the SMR monitoring in Indonesia. Finally, ELC sincerely thanks the Conservation Genetics team at the

Royal (Dick) School of Veterinary Studies for the inspiring working environment and insightful conversations surrounding this research.

## Author contributions

EC and MTPG conceptualised the project. EC carried out lab work, and performed bioinformatic analysis with guidance from GHA. EC drafted the original manuscript and all co-authors contributed edits and approved the final version.

## Availability of data and materials

The raw whole-genome sequencing data for the Singapore bird park’s (Jurong) Magpie-robin (*Copsychus saularis adamsi*) has been deposited to the SRA and is available under BioProject number PRJNA942117.

## References

Andersson, A., Tilley, H., Lau, W., Dudgeon, D., Bonebrake, T., & Dingle, C. (2021). CITES and beyond: Illuminating 20 years of global, legal wildlife trade. Global Ecology and Conservation. doi: 10.1016/j.gecco.2021.e01455.

Adavoudi. R., & Pilot, M. (2021). Consequences of Hybridization in Mammals: A Systematic Review. Genes (Basel). doi: 10.3390/genes13010050.

Broad Institute (2019). “Picard Toolkit.”. GitHub Repository. https://broadinstitute.github.io/picard/;Broad Institute

Burivalova, Z., Lee, T., Hua, F., Lee, J., Prawiradilaga, D., & Wilcove, D. (2017). Understanding consumer preferences and demography in order to reduce the domestic trade in wild-caught birds. Biological Conservation, 209, 423–443. doi: 10.1016/j.biocon.2017.03.005.

Cardoso, P., Amponsah-Mensah, K., Barreiros, J., Bouhuys, J., Cheung, H., Davies, A, Kumschick, S., Longhorn, S., Martínez-Muñoz, C., Morcatty, T., Peters, G., Ripple, W., Rivera-Téllez, E., Stringham, O., Toomes, A., Tricorache, P., & Fukushima, C. (2021). Scientists’ warning to humanity on illegal or unsustainable wildlife trade, Biological Conservation. doi: 10.1016/j.biocon.2021.109341.

Cavill, E.L., Gopalakrishnan, S., Puetz, L.C.,Ribeiro, Â.M., Mak, S.S.T., da Fonseca, R.R., Pacheco, G., Dunlop, B., Accouche, W., Shah, N., Zora, A., Calabrese, L., Genner, M., Jones, G., Guo, C., Zhang, G., & Gilbert, M.T.P. (2022). Conservation genomics of the endangered Seychelles Magpie-Robin (Copsychus sechellarum): a unique insight into the history of a precious endemic bird. Ibis, 164(2), 396–410 doi: 10.1111/ibi.13023

Chng, S.C.L., & Eaton, J.A. (2016). In the Market for Extinction: Eastern and Central Java. TRAFFIC. Petaling Jaya, Selangor, Malaysia. https://www.traffic.org/site/assets/files/2393/in-the-market-for-extinction.pdf. Accessed 13 April 2023.

Chng, S., Shepherd, C., & Eaton, J. (2018). In the market for extinction: birds for sale at selected outlets in Sumatra. TRAFFIC Bulletin. 30(1), 15–22. https://www.traffic.org/site/assets/files/10567/bulletin-30_1-final-web.pdf. Accessed 13 April 2023.

Chng, S.C.L., Saaban, S., Wechit, A., & Krishnasamy, K. (2021). Smuggled For Its Song: The Trade in Malaysia’s Oriental Magpie-robins. TRAFFIC, Southeast Asia Regional Office, Petaling Jaya, Selangor, Malaysia. https://www.traffic.org/site/assets/files/13547/smuggled_for_its_song_oriental_magpie_robin_trade_malaysia_en_report_5_march.pdf. Accessed 13 April 2023.

CITES (2023) Appendices I, II, & III. https://cites.org/sites/default/files/eng/app/2023/E-Appendices-2023-02-23.pdf. Accessed April 13 2023.

Danecek, P., Bonfield, J., Liddle, J., Marshall, J., Ohan, V., Pollard, M., Whitwham, A., Keane, T., McCarthy, S., Davies, R., & Li. H. (2021). Twelve years of SAMtools and BCFtools. GigaScience. doi: 10.1093/gigascience/giab008

Darriba, D., Posada, D., Kozlov, A., Stamatakis, A., Morel, B., & Flouri, T. (2019). ModelTest-NG: A New and Scalable Tool for the Selection of DNA and Protein Evolutionary Models. Molecular biology and evolution, 37(1), 291–294. doi: 10.1093/molbev/msz189.

Eaton, J.A., Leupen, B.T.C., & Krishnasamy, K. (2017). Songsters of Singapore: An Overview of the Bird Species in Singapore Pet Shops. TRAFFIC. Petaling Jaya, Selangor, Malaysia.

Gill, F.B., & Donsker, D. (2014). IOC World Bird List (v4.4). doi: 10.14344/IOC.ML.4.4.

Harris, J.B.C., Green, J.M.H., Prawiradilaga, D.M., Giam, X., Giyanto Hikmatullah, D., Putra, C.A., & Wilcove, D.S. (2015). Using market data and expert opinion to identify overexploited species in the wild bird trade. Biological Conservation, 187, 51–60, doi: 10.1016/j.biocon.2015.04.009

IUCN (2019). Guidelines for the management of confiscated, live organisms. Gland, Switzerland. https://portals.iucn.org/library/sites/library/files/documents/2019-005-En.pdf. Accessed 13 April 2023.

Jones, K.E., Patel, N.G., Levy, M.A., Storeygard, A., Balk, D., Gittleman, & J.L., & Daszak, P. (2008). Global trends in emerging infectious diseases. Nature, 451(7181), 990–993. Doi: 10.1038/nature06536.

Kapp, J.D., Green, R.E., & Shapiro, B. (2021). A Fast and Efficient Single-stranded Genomic Library Preparation Method Optimized for Ancient DNA. Journal of Heredity, 112(3), 241–249. doi: 10.1093/jhered/esab012.

Katoh, K., & Standley, D.M. (2013). MAFFT multiple sequence alignment software version 7: improvements in performance and usability. Molecular Biology and Evolution, 30(4), 772–80. doi: 10.1093/molbev/mst010.

Lee, J.G.H., Chng, S.C.L., & Eaton, J.A. (eds) (2016). Conservation strategy for Southeast Asian songbirds in trade. Recommendations from the first Asian Songbird Trade Crisis Summit 2015 held in Jurong Bird Park, Singapore, 27–29 September 2015. https://www.traffic.org/site/assets/files/2275/conservation-strategy-for-southeastasian-songbirds-in-trade.pdf. Accessed 13th April 2023.

Letunic, I., & Bork, P. (2021). Interactive Tree Of Life (iTOL) v5: an online tool for phylogenetic tree display and annotation, Nucleic Acids Research, 49(W1), W293–W296. doi: 10.1093/nar/gkab301

Leupen, B., Krishnasamy, K., Shepherd, C., Chng, S., Bergin, D., Eaton, J., Yukin, A., Koh, S., Hue, P., Miller, A., Nekaris, K, Nekaris, V., Nijman, S., Saaban Muhammed, A., & Imron, M. (2019). Trade in White-rumped Shamas Kittacincla malabarica demands strong national and international responses. Forktail, 34, 1–8. doi: 10.13140/RG.2.2.17829.04320.

Li, H., & Durbin, R. (2009). Fast and accurate short read alignment with Burrows-Wheeler transform. Bioinformatics, 25(14), 1754–60. doi: 10.1093/bioinformatics/btp324

Lim, H., Zou, F., Taylor, S., Marks, B., Moyle, R., Voelker, G., & Sheldon, F. (2010). Phylogeny of magpie-robins and shamas (Aves: Turdidae: Copsychus and Trichixos): Implications for island biogeography in Southeast Asia. Journal of Biogeography, 37(10), 1894–1906. doi: 10.1111/j.1365-2699.2010.02343.x.

Lim, H., Gawin, D., Shakya, S., Harvey, M., Rahman., M.A., & Sheldon, F. (2017). Sundaland’s east-west rain forest population structure: Variable manifestations in four polytypic bird species examined using RAD-Seq and plumage analyses. Journal of Biogeography, 44(10), 2259–2271. doi: 10.1111/jbi.13031.

Morton, O., Scheffers, B., & Haugaasen, T., & Edwards, D. (2021). Impacts of wildlife trade on terrestrial biodiversity. Nature Ecology & Evolution, 5(4), 1–9. doi:10.1038/s41559-021-01399-y.

Ng, E., Garg, K., Low, G., Chattopadhyay, B., Oh, R., Lee, J., & Rheindt, F. (2017).Conservation genomics identifies impact of trade in a threatened songbird. Biological Conservation, 214, 101–108. doi: 10.1016/j.biocon.2017.08.007.

Okarda, B., Muchlish, U., Kusumadewi, S.D., & Purnomo, H. (2022). Categorizing the songbird market through big data and machine learning in the context of Indonesia’s online market. Global Ecology and Conservation. doi: 10.1016/j.gecco.2022.e02280.

Peng, L., Yang, D., & Lu, C. (2016). Complete mitochondrial genome of oriental magpie-robin Copsychus saularis (Aves: Muscicapidae). Mitochondrial DNA Part B: Resources, 1(1), 21–22. doi: 10.1080/23802359.2015.1137802.

Quinlan, A., & Hall, I. (2010). BEDTools: a flexible suite of utilities for comparing genomic features. Bioinformatics, 26(6), 841–842. doi: 10.1093/bioinformatics/btq033.

Ribeiro, J., Reino, L., Schindler, S., Strubbe, D., Vall-Ilerosa, M., Araújo, M., Carette, M., Mazzoni, S., Monteiro, M., Moreira, F., Rocha, R., Tella, J., Vas, A., Vicente, J., & Nuno, A. (2019). Trends in legal and illegal trade of wild birds: a global assessment based on expert knowledge. Biodiversity Conservation, 28, 3343–3369. doi: 10.1007/s10531-019-01825-5.

Rush, E., Dale, E. & Aguirre A. (2021). Illegal Wildlife Trade and Emerging Infectious Diseases: Pervasive Impacts to Species, Ecosystems and Human Health. Animals. doi: 10.3390/ani11061821.

Scheffers, B., Oliveira, B., Lamb, I., & Edwards, D. (2019). Global wildlife trade across the tree of life. Science, 366(6461), 71–76. doi: 10.1126/science.aav5327

Schubert, M., Ermini, L., Sarkissian, C.D., Jónsson, H., Ginolhac, A., Schaefer, R., Martin M.D., Fernández, R., Kircher, M., McCue, M., Willerslev, E., & Orlando, L. (2014). Characterization of ancient and modern genomes by SNP detection and phylogenomic and metagenomic analysis using PALEOMIX. Nature Protocols, 9, 1056–1082. doi: 10.1038/nprot.2014.063

Schubert, M., Lindgreen, S., & Orlando, L. (2016). AdapterRemoval v2: rapid adapter trimming, identification, and read merging. BMC Research Notes. doi: 10.1186/s13104-016-1900-2

Sheldon, F., Lohman, D., Lim, H., Zou, F., Goodman, S., Prawiradilaga, D., Winker, K., Braile, T., & Moyle, R. (2009). Phylogeography of the magpie-robin species complex (Aves: Turdidae: Copsychus) reveals a Philippine species, an interesting isolating barrier and unusual dispersal patterns in the Indian Ocean and Southeast Asia. Journal of Biogeography, 36(6), 1070–1083. doi: 10.1111/j.1365-2699.2009.02087.x.

Siriwat, P., & Nijman, V. (2020) Wildlife trade shifts from brick-and-mortar markets to virtual marketplaces: A case study of birds of prey trade in Thailand. Journal of AsiaPacific Biodiversity, 13(3), 454–461. doi: 10.1016/j.japb.2020.03.012.

Stamatakis, A. (2014). RAxML version 8: a tool for phylogenetic analysis and post-analysis of large phylogenies. Bioinformatics, 30(9), 1312–1313. doi: 10.1093/bioinformatics/btu033.

Voelker, G., Peñalba, J., Huntley. J., & Bowie, R. (2014). Diversification in an Afro-Asian songbird clade (Erythropygia–Copsychus) reveals founder-event speciation via transoceanic dispersals and a southern to northern colonization pattern in Africa. Molecular Phylogenetics and Evolution, 73, 97–105. doi: 10.1016/j.ympev.2014.01.024.

Wasser, S.K., Joseph Clark, W., Drori, O., Stephen Kisamo, E., Mailand, C., Mutayoba, B., & Stephens, M. (2008). Combating the illegal trade in African elephant ivory with DNA forensics. Conservation Biology, 22(4), 1065–1071. doi: 10.1111/j.1523-1739.2008.01012.x.

Wu, M., Lau, C., Ng, E., Baveja, P., Gwee, C., Sadanandan, K., Ferasyi, T.,Haminuddin & Ramadhan, R., Menner, J., & Rheindt, F. (2022). Genomes From Historic DNA Unveil Massive Hidden Extinction and Terminal Endangerment in a Tropical Asian Songbird Radiation. Molecular Biology and Evolution, 39(9), 1537–1719. doi: 10.1093/molbev/msac189.

WWF (2020) Wildlife crimes are wild crimes against life. https://wwfcee.org/our-offices/slovakia/wildlife-crimes-are-wild-crimes-against-life#:~:text=Wildlife%20crimes%2C%20such%20as%20illegal,to%20species%2C%20after%20habitat%20loss. Accessed 13 April 2023

Wyatt, T., Maher, J., Allen, D., Clarke, N., & Rook, D. (2021). The welfare of wildlife: an interdisciplinary analysis of harm in the legal and illegal wildlife trades and possible ways forward. Crime Law and Social Change, 77, 69–89. doi: 10.1007/s10611-021-09984-9.

